# Adaptive Sampling as tool for Nanopore direct RNA-sequencing

**DOI:** 10.1101/2022.10.14.512223

**Authors:** Isabel S. Naarman-de Vries, Enio Gjerga, Catharina L.A. Gandor, Christoph Dieterich

## Abstract

ONT long-read sequencing provides real-time monitoring and controlling of individual nanopores. Adaptive sampling enriches or depletes specific sequences in Nanopore DNA sequencing, but was not applicable to direct sequencing of RNA so far.

Here, we identify essential parameter settings for direct RNA sequencing (DRS). We demonstrate the superior performance of depletion over enrichment and show that adaptive sampling efficiently depletes specific transcripts in transcriptome-wide sequencing applications. Specifically, we applied our adaptive sampling approach to polyA+ RNA samples from human cardiomyocytes and mouse whole heart tissue. Herein, we show more than 2.5-fold depletion of highly abundant mitochondrial-encoded transcripts that in normal sequencing account for up to 40% of sequenced bases in heart tissue samples.

## Background

Adaptive sampling (AS) describes an add-on technique of the single molecule sequencing technology introduced by Oxford Nanopore Technologies (ONT) [1] to select specific reads during sequencing. The combination of real time basecalling and voltage reversal at specific pores allows to reject molecules that are not of interest. The general approach has been conceptualized by ONT, but the underlying “Read until” scripts were implemented by Loose and coworkers [2]. These initial scripts used ‘squiggle’ data and a dynamic time warping algorithm [2]. GPU basecalling nowadays enables adaptive sampling based on short chunks of basecalled nucleotide sequences [3]. By this approach, individual read sampling strategies can be implemented based on either a black list (depletion mode) or white list (enrichment mode) provided as simple FASTA file. In parallel, several other approaches have been implemented and published to enable Nanopore adaptive sampling. For example, UNCALLED uses raw signal [4], whereas RUBRIC also uses sequencing chunks to filter out unwanted reads [5]. Furthermore, BOSS-RUNS allows dynamic decisions during sequencing in real time [6].

To our best knowledge and despite its relevance in direct RNA-sequencing (DRS) experiments, the feasibility of adaptive sampling for DRS has not been shown yet. This is especially of interest as DRS experiments are usually limited by read number, especially comparing to short read sequencing approaches. Taking advantage of a simple model system composed of two *in vitro* transcripts (IVT), we derive general parameters and characteristics of direct RNA-seq adaptive sampling (DRAS). We show that depletion outcompetes enrichment in terms of efficiency and specificity.

We applied DRAS to specifically deplete mitochondrial derived transcripts (mt-RNA) from direct RNA-seq runs of mouse heart tissue and human induced pluripotent stem cell-derived cardiomyocytes (hiPSC-CM). In libraries from cardiac cells prepared according to standard protocols up to 30-40% of all reads originate from mt-RNAs [7, 8]. These reads are mainly composed of mt-mRNAs and mt-rRNAs that are polyadenylated as well [9]. However, especially for long read sequencing technologies, one is usually more interested in the analysis of cytosolic mRNAs and mt-RNAs may occupy valuable sequencing capacity [10]. We show that DRAS depletes mt-RNAs efficiently and improves detection of transcripts derived from nuclear chromosomes, both for mouse and human samples. As DRAS does not lead to a faster deterioration of active pores, as occasionally observed for sequencing of DNA [11], it represents a simple method to deplete unwanted transcripts especially from samples with limited availability without prior purification that may be associated with sample loss.

## Results and Discussion

### Evaluation of adaptive sampling for direct RNA-seq

Despite the well supported application of AS to Nanopore sequencing of DNA, it has, to our knowledge, not been established for direct sequencing of RNA. In general, two file types can be applied to analyze the adaptive sampling outcome (Figure 1A). On the one hand, the “adaptive sampling” file provides decision information on every sequenced chunk (small read segment). Chunks that are classified as “stop_receiving” are accepted and the read is sequenced completely, whereas “unblock” results in rejection of the read. Chunks that are too short for classification are categorized as “no_decision” (Figure 1A, left panel). On the other hand, the adaptive sampling outcome can be derived from the “sequencing summary” that provides the end reason for every read (Figure 1A, right panel). Reads that are completely sequenced without intervention are classified as “signal_positive", whereas reads rejected in the adaptive sampling process are labeled as “data_service_unblock_mux_change”. Thus, both labels, “unblock” and “data_service_unblock_mux_change", can be used to identify rejected reads.

**Figure 1:**
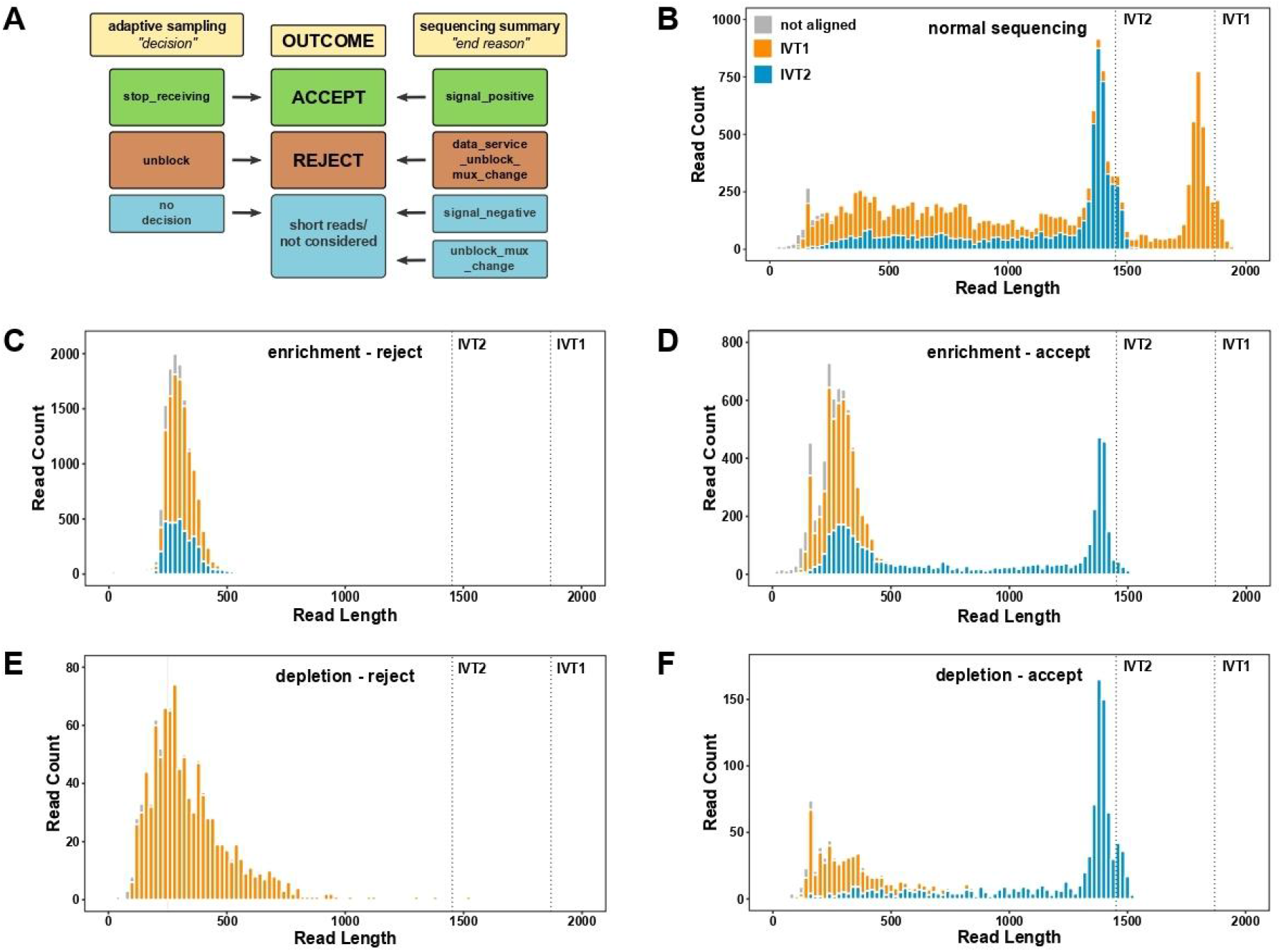
Set up of direct RNA-seq adaptive sampling (DRAS) employing an IVT model system. A) Schematic representation of the classification of reads in adaptive sampling. During the sequencing run two files are generated that can be used to identify accepted and rejected reads, “adaptive sampling” and “sequencing summary”. The “adaptive sampling” file provides information on the decision that was made on individual sequencing chunks. Here “stop_receiving” corresponds to accepted reads, whereas “unblock” represents rejected reads. Very short chunks can not be classified and are categorized as “no_decision”. The “sequencing summary” file provides information on the end reason for every read, which are classified as: “signal_positive” (read passed pore completely), “data_service_unblock_mux_change” (read was rejected during adaptive sampling), “signal_negative” (current delta of 80 pA was observed) and “unblock_mux_change” (strand blocked pore and was rejected). B-F) Equimolar mixtures of IVT1 (1869 nts) and IVT2 (1452 nts) were sequenced on Flongle flow cells in normal sequencing mode (B), adaptive sampling with enrichment of IVT2 (C,D) or adaptive sampling with depletion of IVT1 (E,F). The obtained reads were aligned to the reference sequences and splitted into rejected (C,E) and accepted reads (D,F) based on the sequencing summary.

We determined general properties and characteristics of direct RNA-seq adaptive sampling (DRAS) employing a simple model system of two defined in vitro transcripts (IVT). Equimolar amounts of IVT1 (1869 nt) and IVT2 (1452 nt) were sequenced on Flongle flow cells (Supplementary Figure 1, Figure 1) either in normal sequencing (NS) mode (Figure 1B), AS mode with enrichment of IVT2 (Figure 1C,D) or depletion of IVT1 (Figure 1E,F). The sequencing results were analyzed by the end reason. As expected, the NS setup generates mostly “signal_positive” reads (Supplementary Figure 1A). For the enrichment of IVT2, 60.4% of reads correspond to reads rejected by “data_service_unblock_mux_change” (Supplementary Figure 1B), whereas in depletion mode, 39.6% of reads were rejected (Supplementary Figure 1C). The majority of these reads is shorter than 500 nts, however especially in depletion mode also longer reads are detected (Supplementary Figure 1C).

The alignment of the obtained reads to the IVT1 and IVT2 reference sequences revealed that the expected peaks of the full length transcripts were detected in NS mode (Figure 1B). The aligned data of the adaptive sampling approaches were splitted into accepted and rejected reads according to the sequencing summary. Here, full length IVT1 was not detectable either upon enrichment of IVT2 (Figure 1C,D) or depletion of IVT1 (Figure 1E,F). The majority of the IVT1 reads corresponds to the reads rejected by AS (Figure 1C,E). However, we noticed also a substantial fraction of IVT2 reads below 500 nts that are rejected in enrichment mode (Figure 1C). This coincides with a larger fraction of erroneously accepted IVT1 reads in enrichment (Figure 1D) compared to depletion (Figure 1F). Based on the number of sequenced bases per transcript (Supplementary Figure 1D-F), we calculated an enrichment factor (Supplementary Information) of 1.35 for the enrichment mode and 1.75 for the depletion mode, indicating a higher efficiency of depletion in the context of direct RNA-seq experiments. The enrichment factor of 1.75 is close to the theoretically possible enrichment factor of 1.86, as calculated according to Martin et al. [12] (Supplementary Information). Furthermore, Calculation of the error rates (Supplementary Information) revealed a higher specificity and efficiency of the depletion mode, with no wrongly rejected reads and 32% wrongly accepted reads, compared to 42% wrongly accepted and 28% wrongly rejected reads in the enrichment mode.

### Application of adaptive sampling to depletion of mitochondrial transcripts from mouse heart

Based on the findings we tested a “real life” DRAS use case and choose the depletion of mitochondrial encoded RNAs (mt-RNAs). mt-RNAs that are highly expressed in heart tissue [7, 8] can disturb the detection of cytosolic transcripts due to the limited sequencing capacity available in DRS experiments. We recently introduced a method for the experimental depletion of mitochondrial transcripts by RNase H cleavage, mt-clipping [10]. Here, we present DRAS as alternative strategy to deplete mt-RNAs without prior purification. We applied this method to mouse heart tissue (Figure 2, Supplemental Figure 2-4) as well as to hiPSC-CM (Supplementary Figures 5-7). The depletion of mt-RNAs was analyzed in duplicate samples on MinION flow cells, from which 50% of pores were used for NS and 50% for AS. Obtained reads were splitted into accepted (“stop receiving”) and rejected (“unblock”) reads, which should represent cytoplasmic and mitochondrial RNAs, respectively (Figure 2A). As observed for the IVT model, most AS rejected reads are shorter than 500 nts (Supplementary Figure 2A-D). Compared to the normal sequencing, the total number of mitochondrial reads in adaptive sampling was reduced from 32.2% to 22.5%, including the rejected reads which are usually very short (Figure 2B,C). Strikingly, the number of sequenced bases mapping to the mitochondrial chromosome were drastically reduced in adaptive sampling from 42% to 16.5%. Importantly, no mt-RNAs were wrongly accepted and the rejected reads were composed mainly of short reads (Figure 2C, Supplementary Figure 3A). Reads mapping to the other chromosomes were mainly increased (Figure 2B, Supplementary Figure 3A), with the exception of chromosome 1, where many mitochondrial pseudogenes are located in mice [13]. In total, 99% of rejected reads mapped to the mitochondrial chromosome and chromosome 1 (Supplementary Table 1). To exclude a potential detrimental effect on sequencing output due to faster detoriation of pores, we analyzed pore health separately for NS and AS and found no differences between sequencing conditions (Supplementary Figure 2E-H). Exemplarily, we analyzed detection of genes implicated in dilated cardiomyopathy (DCM). DRAS enhanced detection of many genes of the gene set “HP_DILATED_CARDIOMYOPATHY” (Supplementary Figure 4). Exemplarily, we show here a higher coverage along the gene body for Gatad1 [14] (Figure 2D) and longer reads for the junction plakoglobin Jup [15] (Figure 2E).

**Figure 2:**
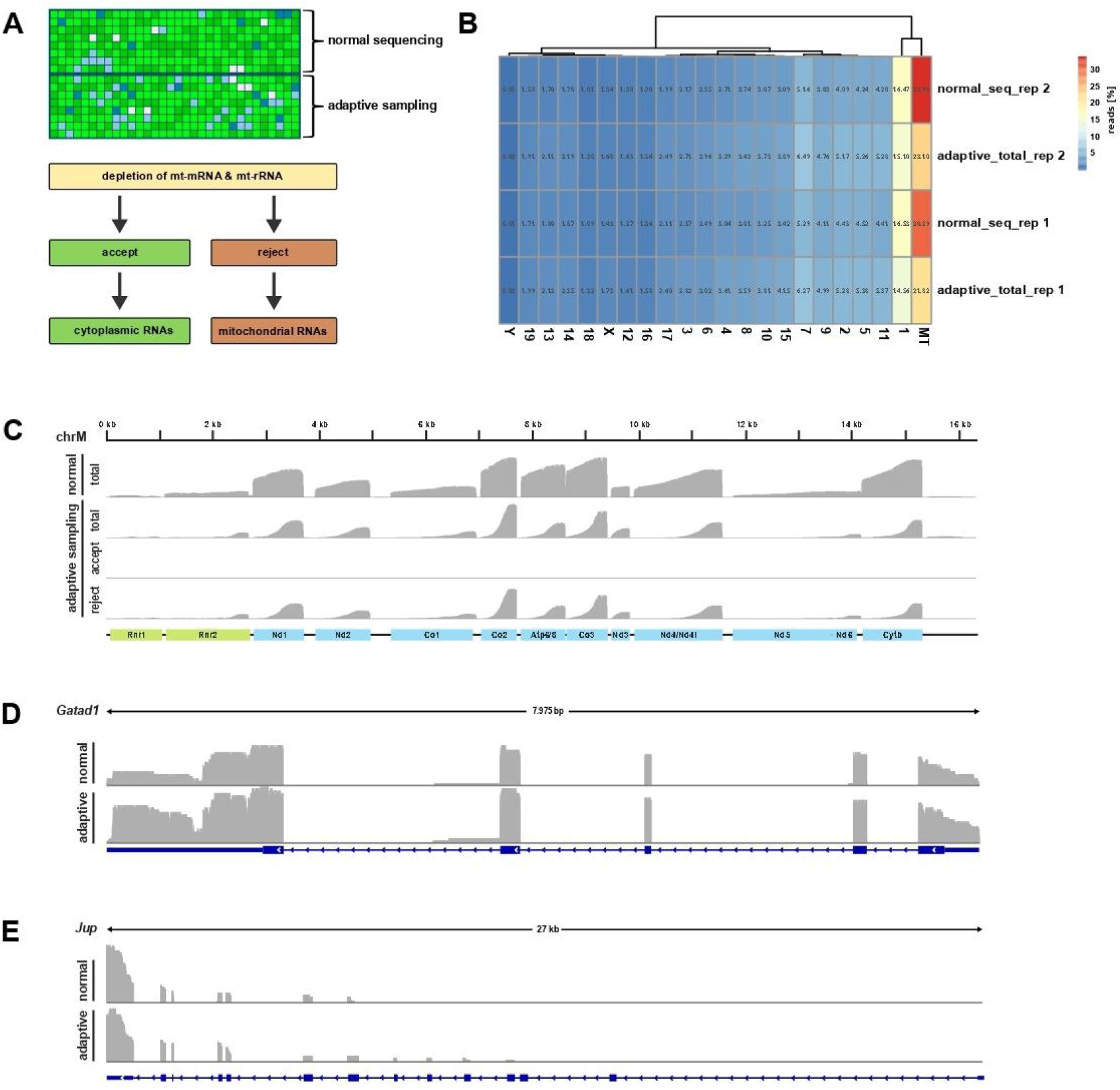
Depletion of mt-RNAs in direct RNA-seq of mouse heart tissue samples. A) Schematic representation of the experimental setup. Upper panel: Libraries were sequenced on a MinION flow cell, with 50% percent of channels in normal sequencing mode and 50% of channels in adaptive sampling mode. Lower panel: Reads were splitted prior to further analysis according to the adaptive sampling decision. B) Gene counts per chromosome for two biological replicates splitted into “normal sequencing” and “total reads from adaptive sampling”. C) Normalized coverage of reads mapped to the mitochondrial chromosome (chrM) in the individual samples as indicated. D,E) Normalized coverage of Gatad1 (D) and Jup (E) in normal sequencing and adaptive sampling as indicated.

Addtionally, we applied the depletion of mitochondrial transcripts by DRAS to human iPSC-CM that were employed for the mt-clipping protocol [10] as well. General characteristics as read length histogram and pore health were highly comparable to the mouse experiment (Supplementary Figure 5). Compared to the primary mouse tissue, the fraction of mt-RNAs is smaller (Supplementary Figure 6A, 9.84% of reads in normal sequencing), whereas the efficiency of depletion is highly comparable. In hiPSC-CM the number of sequenced bases mapping to the mitochondrial chromosome is reduced 2.8-fold from 6.7% to 2.4% by adaptive sampling. The depletion of mt-RNAs (Supplementary Figure 6A-C) is accompanied by an increased number of reads mapped to the nuclear chromosomes (Supplementary Figure 6A). In contrast to mouse, we do not observe a co-depletion of mitochondrial pseudogenes for the human samples and 98% of rejected reads mapped to the mitochondrial chromosome (Supplementary Table 2). Similar to the mouse samples, almost no full length transcripts of mt-RNAs were detected in the AS reads (Supplementary Figure 6D,E). To compare the mt-RNA depletion by AS with the experimental depletion by mt-clipping [10], we analyzed detection of genes implicated in pathogenesis of DCM [16]. The obtained number of reads per gene is increased for most DCM related genes in AS over NS in both replicates (Supplementary Figure 7A), as illustrated for MYH6 (Supplementary Figure 7B) [17] and PLN [18] (Supplementary Figure 7C).

## Conclusions

In this work, we show for the first time that the AS option provided by ONT is suitable for direct RNA-seq as well and may be benefical in several experimental setups and biological problems.

Taking advantage of a simple model system composed of two defined *in vitro* transcripts, we determine essential parameters of DRAS. We show that decisions are made based on sequencing chunks that are shorter than for sequencing of DNA, mainly due to the slower translocation speed of RNA compared to DNA, rendering it usefull also for the targeting of relatively short transcripts. Furthermore, we provide evidence that the depletion of specific transcripts has a higher efficiency and specificity than the enrichment.

As illustrated in Figure 1A, AS may be analyzed either on the “adaptive sampling” or the “sequencing summary” file. Although in theory the reads classified as “unblock” or “data_service_unblock_mux_change” should be identicial, we observed minor differences in classification between the two files. Therefor, we recommend to use only one file system throughout analysis.

As use case in a vertebrate transcriptome-wide experimental setup, we established the depletion of mt-RNAs from heart samples (mouse heart tissue and hiPSC-CM). Mt-RNAs are highly enriched in cardiac cells, as a high number of mitochondria is required to meet the high oxygen demand of these cells. Because these transcripts can make up to 30-40% of the polyadenylated RNA fraction [7, 8], and may not be of general interest, they block valuable sequencing capacity for cytosolic mRNAs. We show here that, despite the short nature of the mt-RNAs, AS for depletion of mt-RNAs is beneficial for detection of cytosolic mRNAs, exemplified for a group of genes implicated in DCM.

Although the depletion mode applied here has a higher efficiency in direct RNA- seq, also the enrichment of specific transcripts may be beneficial, e.g. if one is interested in the in depth knowledge on isoform expression of a specific subgroup of genes. This may be of relevance as well in the understanding of cardiovascular diseases, as many disease entities, as cardiomyopathies, are caused by aberrant splicing regulation [19].

## Methods

### Isolation of total RNA from hiPSC-CM and mice samples

Total RNA from hiPSC-CM (generated as described in [10]) was isolated using Trizol (Thermo Fisher Scientific) as described previously [10]. Mice heart tissue sections from Sham operated mice were homogenized in 750 μl Qiazol (Qiagen) using the TissueLyser (Qiagen) (4-5 times, 1 min, 30 Hz). Afterwards, RNA was isolated with the miRNeasy Mini kit (Qiagen) according to the manufacturers protocol.

### Isolation of polyA + RNA

For isolation of polyA + RNA from hiPSC-CM total RNA, 30 μg total RNA was digested with 1 μl DNase I (New England Biolabs) in a total volume of 100 μl for 10 min at 37°C followed by inactivation of the enzyme (10 min, 65 ° C). 50 μl Dynabeads Oligo(dT) 25 beads were washed once with binding buffer (10 mM Tris pH 7.5, 1M lithium chloride, 6.5 mM EDTA) and resuspended in 110 μl binding buffer. Beads and DNase I-treated RNA were combined and incubated 5 min at room temperature. Beads were collected on a magnet and washed two times with 200 μl washing buffer (5 mM Tris pH 7.5, 150 mM lithium chloride, 1 mM EDTA). RNA was eluted in 100 μl by heating to 70°C for 2 min. Beads were resuspended in 100 μl binding buffer and recombined with the eluted RNA for a second purification round as described above. Final polyA^+^ RNA was eluted in a volume of 10 μl. PolyA^+^ RNA from mouse total RNA was isolated with the Oligotex mRNA purification kit (Qiagen) according to the manufacturers protcocol. Concentration and purity of the isolated polyA^+^ RNA was analyzed on a Nanodrop (Thermo Fisher Scientific) and Fragment Analyzer (Agilent), respectively.

### Generation of in vitro transcripts

Generation of the in vitro transcripts has been described previously [20]. IVT1 represents the complete sequence of the human 18S rRNA, whereas IVT2 represents a 3’ fragment of the human 28S rRNA. Both IVTs have been generated with MEGAscript T7 Transcription kit (Thermo Fisher Scientific) according to the manufacturers protocol and purified with Zymo RNA Clean & concentrator kits (Zymo Research).

### Generation of direct RNA-seq libraries

Libraries were generated with 500 ng polyA + RNA per reaction for MinION flow cells and 200 ng RNA equimolar IVT mixture for Flongle flow cells using the direct RNA sequencing kit (SQK-RNA002, Oxford Nanopore Technologies) including the reverse transcription step. Concentration of libraries was determined with the Qubit dsDNA HS kit (Thermo Fisher Scientific). Libraries were loaded completely on a MinION R9.4.1 flow cell (FLO-MIN106D) or Flongle flow cell and sequenced for 24 to 48 hours as outlined below.

### Set up of GPU basecalling and adaptive sampling

The RNA sequencing data were acquired by using MinKNOW v21.10.4 installed on a HP zBook Create G7 Notebook running on Ubuntu 20.04.2 LTS, Intel(R) Core (TM) i7-10850H CPU @ 2.70GHz (1 CPU socket, 6 cores, 12 threads). Live basecalling of the FAST5 files was performed using Guppy 6.0.1 using a built-in NVIDIA GPU (GeForce RTX 2070 Mobile, CUDA version 11.4). Sequencing runs were started via the MinKNOW GUI and SQK-RNA002 kit as well as the respective flow cell chosen. Adaptive sampling was activated either in depletion or enrichment mode and required a FASTA file providing the reference sequences that should be enriched (whitelist) or depleted (blacklist), respectively. For mt-RNA depletion experiments, the range for adaptive sampling was defined to 50% of channels of the MinION flow cell. Guppy was run in high accuracy mode with a Q score threshold of 7.

### Data processing

Subsequent FASTQ read alignment was performed with minimap2 (2.22 -r1101) using the following command options for Nanopore direct RNA sequencing: -ax splice -uf -k14 -secondary=no. Secondary reads were subsequently filtered using the following SAM flags: samtools view -S -b -F 2304. Gene counts were obtained from the generated reads by using featureCounts() function from the Rsubread R-package (GRCh38-105 and GRCm30-105 annotations were used for humans and mouse respectively). For the analysis of the IVT model, the following options were used: minimap2 -ax splice -uf -k14 ref.fa direct-rna.fq > aln.sam. Bam-file analysis and allocation of IVT sequencing data were performed using samtools (1.10.2-3). Read IDs of rejected reads were identified by the sequencing summary file automatically generated after sequencing by filtering for “Data_Service_Unblock_Mux_Change”.

### Visualization of data

Data analysis and visualization of read length histograms were performed in R. Capabilities of base R were expanded by several packages including the tidyverse (1.3.1.) collection (dplyr, tidyr, ggplot). Heatmaps were generated using pheatmap (1.0.12). Coverage tracks were visualized with Integrative Genomics Viewer and schematic figures were drawn with Inkscape.

## Supplementary Information

Supplementary Information/Figures/Tables are available online.

## Acknowledgements

We thank Harald Wilhelmi for excellent support in setting up the computational framework and Jessica Eschenbach for excellent technical assistance. We are grateful to Matthias Dewenter and Johannes Backs for providing the mouse heart tissue samples.

## Abbreviations

AS: Adaptive sampling
NS: Normal sequencing
ONT: Oxford Nanopore Technologies
DRAS: direct RNA-seq adaptive sampling
IVT: in vitro transcript
hiPSC-CM: human induced pluripotent stem cell-derived cardiomyocytes
DCM: dilated cardiomyopathy

## Availability of data and materials

Workflow and sequencing data are provided at Zenodo: https://doi.org/10.5281/zenodo.6581449.

## Authors’ contributions

ISNdV and CLAG performed all experiments. EG and CLAG analyzed direct RNA-seq data. ISNdV, EG and CLAG compiled figures. ISNdV wrote the manuscript. ISNdV and CD designed and supervised the project. All authors critically read the manuscript, revised it and approved the final version.

## Additional Figures

Supplementary Figure 1

Read length histogram analysis corresponding to Figure 1.

Supplementary Figure 2

Read length histogram and pore health analysis for sequencing of mouse samples.

Supplementary Figure 3

Mt-RNA read length analysis of mouse samples.

Supplementary Figure 4

Log2fold change of the normalized gene count of genes in the gene set HP_DILATED_CARDIOMYOPATHY for mouse.

Supplementary Figure 5

Read length histogram and pore health analysis for sequencing of human samples.

Supplementary Figure 6

Analysis of human samples.

