## Supplementary Materials for "Adaptive Sampling as tool for Nanopore direct RNA-sequencing"

### Supplementary Information

#### Supplementary Methods

##### *Calculation of Potential Enrichment for the IVT model*

To calculate the potential enrichment, the mathematical formula of Martin *et al.* (2022) was used: Since the potential enrichment achievable through adaptive sampling depends on several factors such as species abundance, sequencing speed, decision time, and capture time, it is advisable to predict the enrichment prior to the experiment and check if adaptive sampling is a suitable strategy. For the calculation of the possible enrichment the mathematical model of Martin *et al.* (2022) was used and slightly adapted for RNA sequencing. The enrichment factor (E) can be calculated by the obtained abundance of the target species (bases sequenced) by adaptive sampling ($x_{as}$) and normal sequencing ($x_{norm}$): $E = \frac{x_{as}}{x_{norm}}$.

Assuming that the sequencing rate (S) remains nearly constant during the time most reads are sequenced, the enrichment factor can be determined by the quotient of the time that is spend on sequencing the target with adaptive sampling ($T_{AS}$) and without adaptive sampling ($T_{NS}$): $E = \frac{T_{AS}}{T_{NS}}$.

The time that is spend on the target sequence without adaptive sampling can be calculated by the proportion of target molecules (y), the average read length of the target (R), the sequencing rate (S) and time spend on capturing a new molecule (C):

$$T_{NS} = \frac{y\,\frac{R}{S}}{\frac{R}{S} + C} = \frac{yR}{R + CS} \quad (1)$$

Using adaptive sampling, off-target reads are sequenced until the decision is made to eject the read. This decision time (D) for the proportion of off-target reads (1-y) leads to the following formular for $T_{AS}$:

$$T_{AS} = \frac{y\,\frac{R}{S}}{y\frac{R}{S} + (1-y)D + C} = \frac{yR}{yR + (1-y)DS + CS} \quad (2)$$

Taken together formular (1) and (2), the enrichment factor can be calculated by:

$$E = \frac{T_{AS}}{T_{NS}} = \frac{R + CS}{yR + (1-y)DS + CS} \quad (3)$$

In this experiment, the average read length (R) was 1660 bases, and the target species abundance (y) was 50%. Since RNA has a slower sequencing rate than DNA, the given sequencing rate (S) of ~240 bases per second had to be adapted. Previous direct RNA experiments exhibited a sequencing rate of ~ 80 b/s. The capture time (C) of 0.5 s given in the nanopore adaptive sampling sheet was used. Since the time to decide (D) of 1 s refers to DNA sequencing ($\sim\frac{240\ bases}{240\ b/s} = 1\ s$), the decision time was adapted to RNA sequencing. Assuming it takes on average 300 bases until a decision can be made (shown by previous experiments) and RNA gets sequenced with a sequencing rate of ~ 80 b/s, it results in a decision time of ~ 3.75 s. Taking these numbers for the calculation of the formula we get a potential enrichment of 1.66.

$$E = \frac{R + CS}{yR + (1-y)DS + CS} = \frac{1660\,b + 0.5\,s * \,80\,b/s}{0.5 * 1660\,b + (1-0.5) * 3.75\,s * 80\frac{b}{s} + 0.5 * 80\,b/s} = 1.66$$

Since here, unlike in most experiments, the length of the target and non-target reads is known, the formula for $T_{NS}$ can be further specified by introducing the time spend on sequencing an unwanted read (O), in this case IVT1. It is given by the read length ($R_{off}$) divided by (S). This results in the following formula:

$$E = \frac{yR_{On} + (1-y)OS + CS}{yR_{On} + (1-y)DS + CS} = \frac{yR_{On} + (1-y)R_{Off} + CS}{yR_{On} + (1-y)DS + CS} \quad (4)$$

$$= \frac{0.5 * 1452\,b + 0.5 * 1869\,b + 0.5\,s * 80\,b/s}{0.5 * 1452\,b + 0.5 * 3.75\,s * 80\,b/s + 0.5\,s * 80\,b/s} = 1.86$$

Under the assumptions described above, the maximum achievable enrichment by composition in this experiment is 1.86.

#### *Calculation of the observed Enrichment factor for the IVT model*

The enrichment factor (E) was calculated by dividing the proportion of IVT2 bases using adaptive sampling ($x_{AS}$) by the proportion of IVT2 bases using normal sequencing ($x_{NS}$). As shown below, enriching the IVT2 by depleting IVT1 resulted in a higher enrichment factor ($E_{Dep}$ = 1.75) compared to the experiment where the enrichment mode was set ($E_{Enr}$ = 1.35).

| | *Target bases (total) [%]* | *Enrichment Factor (E)* |
|---|---|---|
| *Normal Sequencing* | 40 | 1 |
| *Enrichment* | 54 | 1.35 |
| *Depletion* | 70 | 1.75 |

#### *Calculation of the error rate for the IVT model*

To further investigate the proportion of IVT2 reads that were mistakenly rejected by adaptive sampling, data was divided by "end reason" into two groups. Reads with "data_service_unblock_mux_change" and all remaining reasons were classified as rejected reads and accepted reads, respectively, using the information from the sequencing summary file.

To evaluate the error rates for the corresponding mode, the percentage of IVT2 reads within the rejected reads (wrongly rejected reads) and the percentage of IVT1 reads within the accepted data (wrongly accepted reads) were calculated.

| | *Wrongly rejected reads [%]* | *Wrongly accepted reads [%]* |
|---|---|---|

| | | |
|---|---|---|
| *Enrichment* | 28 | 42 |
| *Depletion* | 0 | 32 |

#### ***Supplementary Figure Legends***

##### ***Supplementary Figure 1:***

Set up of direct RNA-seq adaptive sampling (DRAS) employing an IVT model system. Equimolar mixtures of IVT1 (1869 nts) and IVT2 (1452 nts) were sequenced on Flongle flow cells in normal sequencing mode (A,D), adaptive sampling with enrichment of IVT2 (B,E) or adaptive sampling with depletion of IVT1 (C,F). The obtained reads (A-C) or sequenced bases (D-F) were splitted according to the end reason reported in the sequencing. Reported end reasons were: "signal_positive" (read passed pore completely), "data_service_unblock_mux_change" (read was rejected during adaptive sampling), "signal_negative" (current delta of 80 pA was observed) and "unblock_mux_change" (strand blocked pore and was rejected).

***Supplementary Figure 2:***

Read length histogram and pore health analysis for sequencing of mouse samples. A-D) Read length histogram of normal sequencing (A,C) or depletion of mitochondrial RNAs by adaptive sampling (B,D) of polyA$^+$ RNA derived from mouse whole heart tissue. End reasons are indicated as in Supplementary Figure 1. E-H) Pore health analysis of the sequencing runs as indicated in A-D. The relative fraction of pores in the indicated states were derived from the mux_scan_data and splitted in normal sequencing and adaptive sampling according to the pore number.

***Supplementary Figure 3:***

Mt-RNA read length analysis of mouse samples. A) Normalized coverage of reads mapped to the mitochondrial chromosome (chrM) in the individual samples as indicated for replicate 2. B,C) Length of reads mapping to the individual mitochondrial transcripts in adaptive sampling and normal sequencing of mouse replicate 1 (B) and mouse replicate 2 (C).

***Supplementary Figure 4:***

Log2fold change of the normalized gene count of genes in the gene set HP_DILATED_CARDIOMYOPATHY for the mouse analysis. Genes were filtered for expression using the filterByExpr() function from the edgeR package with the default settings.

***Supplementary Figure 5:***

Read length histogram and pore health analysis for sequencing of human samples. A-D) Read length histogram of normal sequencing (A,C) or depletion of mitochondrial RNAs by adaptive sampling (B,D) of polyA$^+$ RNA derived from hiPSC-derived cardiomyocytes. End reasons are indicated as in Supplementary Figure 1. E-H) Pore health analysis of the sequencing runs as indicated in A-D. The relative fraction of pores in the indicated states were derived from the mux_scan_data and splitted in normal sequencing and adaptive sampling according to the pore number.

***Supplementary Figure 6:***

Analysis of depletion of mitochondrial transcripts from human samples. A) Gene counts per chromosome for two biological replicates splitted into "normal sequencing" and "total reads from adaptive sampling". B,C) Normalized coverage of reads mapped to the mitochondrial chromosome (chrM) in the individual samples of replicate 1 (B) and replicate 2 (C). D,E) Length of reads mapping to the individual mitochondrial transcripts in adaptive sampling and normal sequencing of human replicate 1 (D) and mouse replicate 2 (E).

***Supplementary Figure 7:***

Analysis of DCM related genes in human samples. A) Log2fold change of the normalized gene count of genes implicated in DCM pathogenesis according to Jordan *et al.*, 2021. B,C) Normalized coverage of *MYH6* (B) and *PLN* (C) in normal sequencing and adaptive sampling as indicated.

#### ***Supplementary References***

Jordan E, Peterson L, Ai T, Asatryan B, Bronicki L, Brown E, et al. Evidence-based assessment of genes in dilated cardiomyopathy. Circulation. 2021;144(1):7-19.

Martin S, Heavens D, Lan Y, Horsfield S, Clark MD, Leggett RM. Nanopore adaptive sampling: a tool for enrichment of low abundance species in metagenomic samples. Genome Biology. 2022;23(1):1-27.

### Figure S1

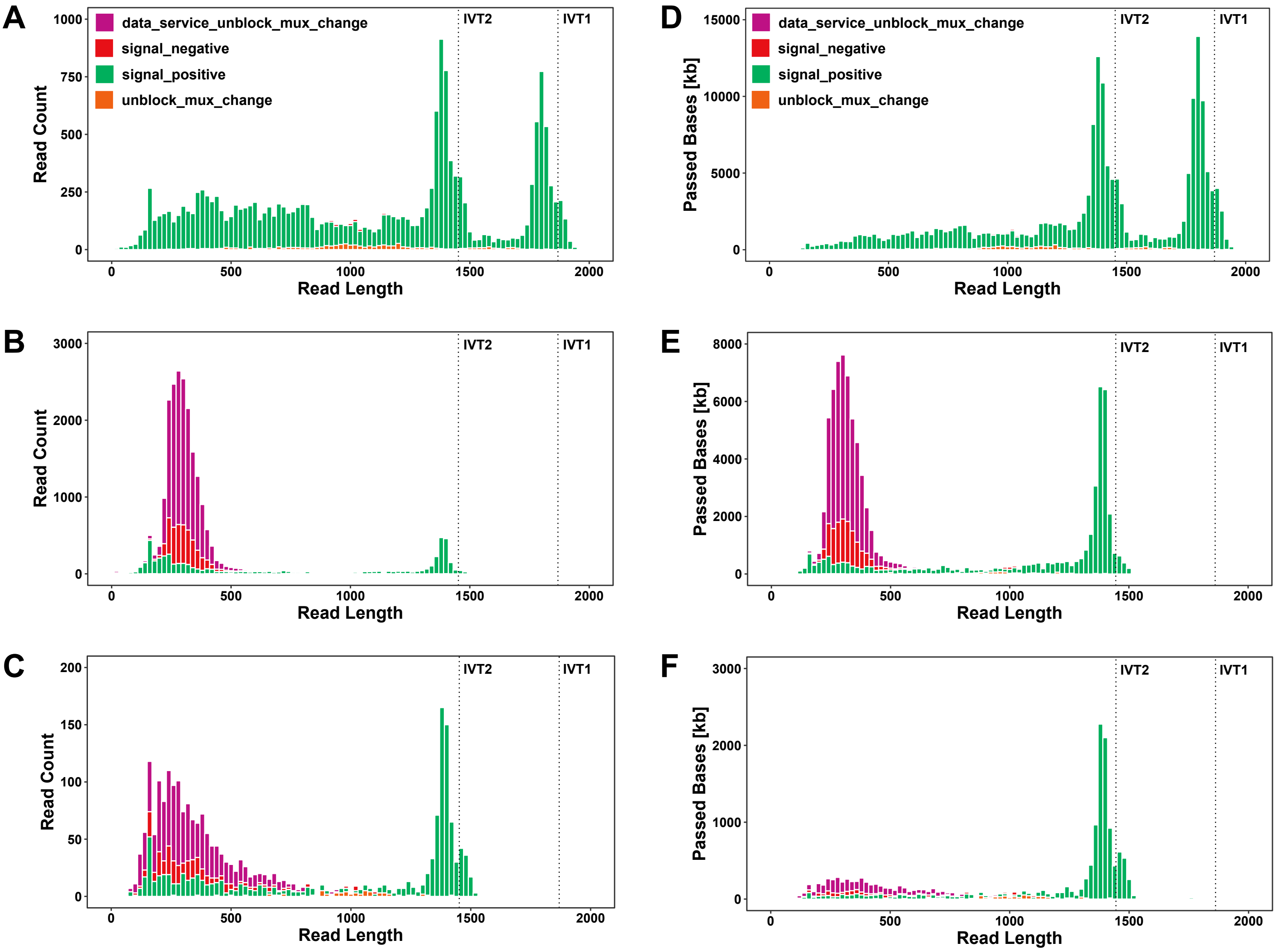
A
B
C
D
E
F
data_service_unblock_mux_change
signal_negative
signal_positive
unblock_mux_change
IVT2
IVT1
Read Count
Passed Bases [kb]
Read Length
0
500
1000
1500
2000
250
750
3000
5000
10000
15000
4000
6000
8000
50
100
150
200

### Figure S2

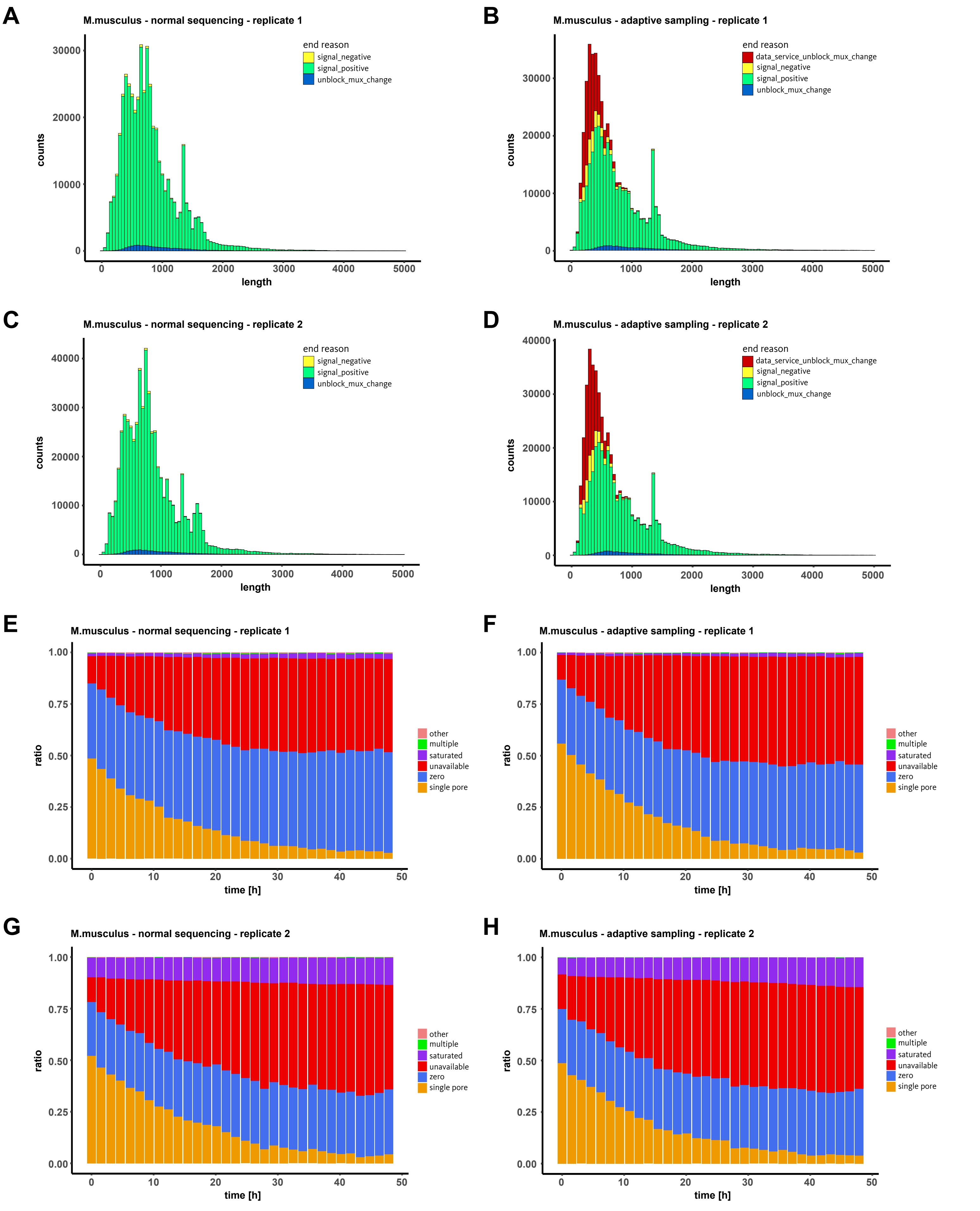

A
M.musculus - normal sequencing - replicate 1
end reason
signal_negative
signal_positive
unblock_mux_change
counts
length
B
M.musculus - adaptive sampling - replicate 1
end reason
data_service_unblock_mux_change
signal_negative
signal_positive
unblock_mux_change
counts
length
C
M.musculus - normal sequencing - replicate 2
end reason
signal_negative
signal_positive
unblock_mux_change
counts
length
D
M.musculus - adaptive sampling - replicate 2
end reason
data_service_unblock_mux_change
signal_negative
signal_positive
unblock_mux_change
counts
length
E
M.musculus - normal sequencing - replicate 1
other
multiple
saturated
unavailable
zero
single pore
ratio
time [h]
F
M.musculus - adaptive sampling - replicate 1
other
multiple
saturated
unavailable
zero
single pore
ratio
time [h]
G
M.musculus - normal sequencing - replicate 2
other
multiple
saturated
unavailable
zero
single pore
ratio
time [h]
H
M.musculus - adaptive sampling - replicate 2
other
multiple
saturated
unavailable
zero
single pore
ratio
time [h]

### Figure S3

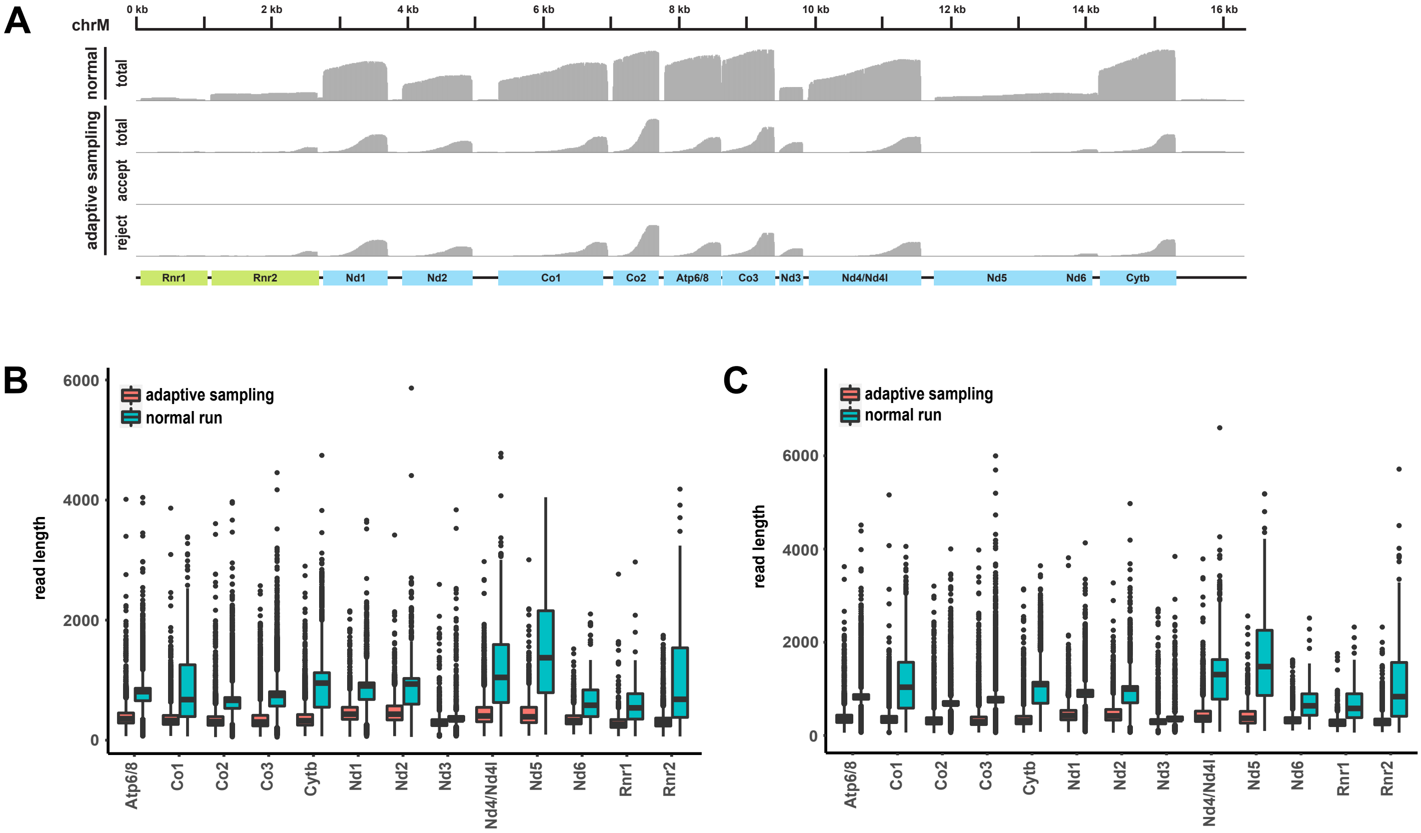

A
chrM
0 kb
2 kb
4 kb
6 kb
8 kb
10 kb
12 kb
14 kb
16 kb
normal
total
adaptive sampling
total
accept
reject
Rnr1
Rnr2
Nd1
Nd2
Co1
Co2
Atp6/8
Co3
Nd3
Nd4/Nd4l
Nd5
Nd6
Cytb
B
adaptive sampling
normal run
read length
6000
4000
2000
0
Atp6/8
Co1
Co2
Co3
Cytb
Nd1
Nd2
Nd3
Nd4/Nd4l
Nd5
Nd6
Rnr1
Rnr2
C
adaptive sampling
normal run
read length
6000
4000
2000
0
Atp6/8
Co1
Co2
Co3
Cytb
Nd1
Nd2
Nd3
Nd4/Nd4l
Nd5
Nd6
Rnr1
Rnr2

### Figure S4

#### Normalized Gene Counts per Bases per Million

| Gene | rep2 | rep1 |
|---|---|---|
| Abcc9 | 0.65 | 0.96 |
| Acad8 | 1.13 | -0.29 |
| Acad9 | -0.21 | 0.56 |
| Acta1 | 0.97 | 0.59 |
| Actc1 | 0.66 | 0.37 |
| Actn2 | 0.35 | 0.39 |
| Ankrd1 | 0.87 | 0.44 |
| Bag3 | 0.28 | 0.67 |
| Bola3 | 0.88 | 0.73 |
| Brcc3 | 0.18 | 0.34 |
| Cap2 | 0.79 | 0.52 |
| Casq2 | 0.57 | 0.44 |
| Cox7b | 0.61 | 0.35 |
| Cpt2 | 0.77 | 0.15 |
| Cryab | 0.52 | 0.36 |
| Csrp3 | 0.52 | 0.39 |
| Des | 0.69 | 0.39 |
| Dmd | 0.84 | 0.34 |
| Dmpk | 0.66 | 0.24 |
| Dnajc19 | 0.41 | 0.00 |
| Dpm3 | 0.41 | -1.08 |
| Dsp | 0.11 | 0.54 |
| Fhl1 | 0.83 | 0.47 |
| Fhl2 | 0.70 | 0.41 |
| Gatad1 | 0.88 | 1.06 |
| Hadh | 0.70 | 0.40 |
| Hadha | 0.83 | 0.55 |
| Hadhb | 0.40 | 0.45 |
| Hand2 | 0.71 | 0.69 |
| Hccs | 0.40 | -0.13 |
| Hmgcl | 0.08 | 0.56 |
| Hspg2 | 0.60 | 0.36 |
| Jph2 | 0.38 | 0.75 |
| Jup | 0.42 | 0.19 |
| Kcnj5 | -0.04 | 1.61 |
| Lamp2 | 0.60 | 0.96 |
| Ldb3 | 0.64 | 0.44 |
| Lims2 | 0.60 | -0.95 |
| Lmna | 0.24 | 0.92 |
| Mlycd | 0.91 | 0.06 |
| Mmachc | 0.69 | 0.44 |
| Mybpc3 | 0.67 | 0.42 |
| Myh6 | 0.34 | 0.20 |
| Myh7 | -0.40 | 0.07 |
| Mypn | -0.09 | -0.44 |
| Ndufaf3 | -0.10 | 1.08 |
| Ndufb11 | 0.57 | 0.58 |
| Ndufb8 | 0.73 | 0.49 |
| Ndufs2 | 0.45 | 0.46 |
| Nexn | 0.46 | 0.74 |
| Pgm1 | 0.47 | 0.80 |
| Pln | 0.71 | 0.43 |
| Rbm20 | 0.30 | 0.93 |
| Rpl3l | 0.58 | 0.37 |
| Ryr2 | 0.09 | 0.28 |
| Scn5a | 0.60 | 1.30 |
| Sdha | 0.63 | 0.54 |
| Sdhd | 0.80 | 0.39 |
| Sgcb | 0.77 | 0.91 |
| Ski | -0.99 | 0.38 |
| Slc25a4 | 0.56 | 0.48 |
| Surf1 | 1.30 | -0.30 |
| Tcap | 0.53 | 0.64 |
| Tnnc1 | 0.52 | 0.54 |
| Tnni3 | 0.55 | 0.50 |
| Tnni3k | 0.33 | 0.58 |
| Tnnt2 | 0.50 | 0.45 |
| Tpm1 | 0.57 | 0.43 |
| Tpm3 | -0.04 | -0.06 |
| Trdn | 0.85 | 0.49 |
| Tsfm | -0.52 | 0.83 |
| Ttn | 0.57 | 0.44 |
| Txnrd2 | 1.47 | 0.90 |

Color scale: 1.5, 1, 0.5, 0, -0.5, -1

### Figure S5

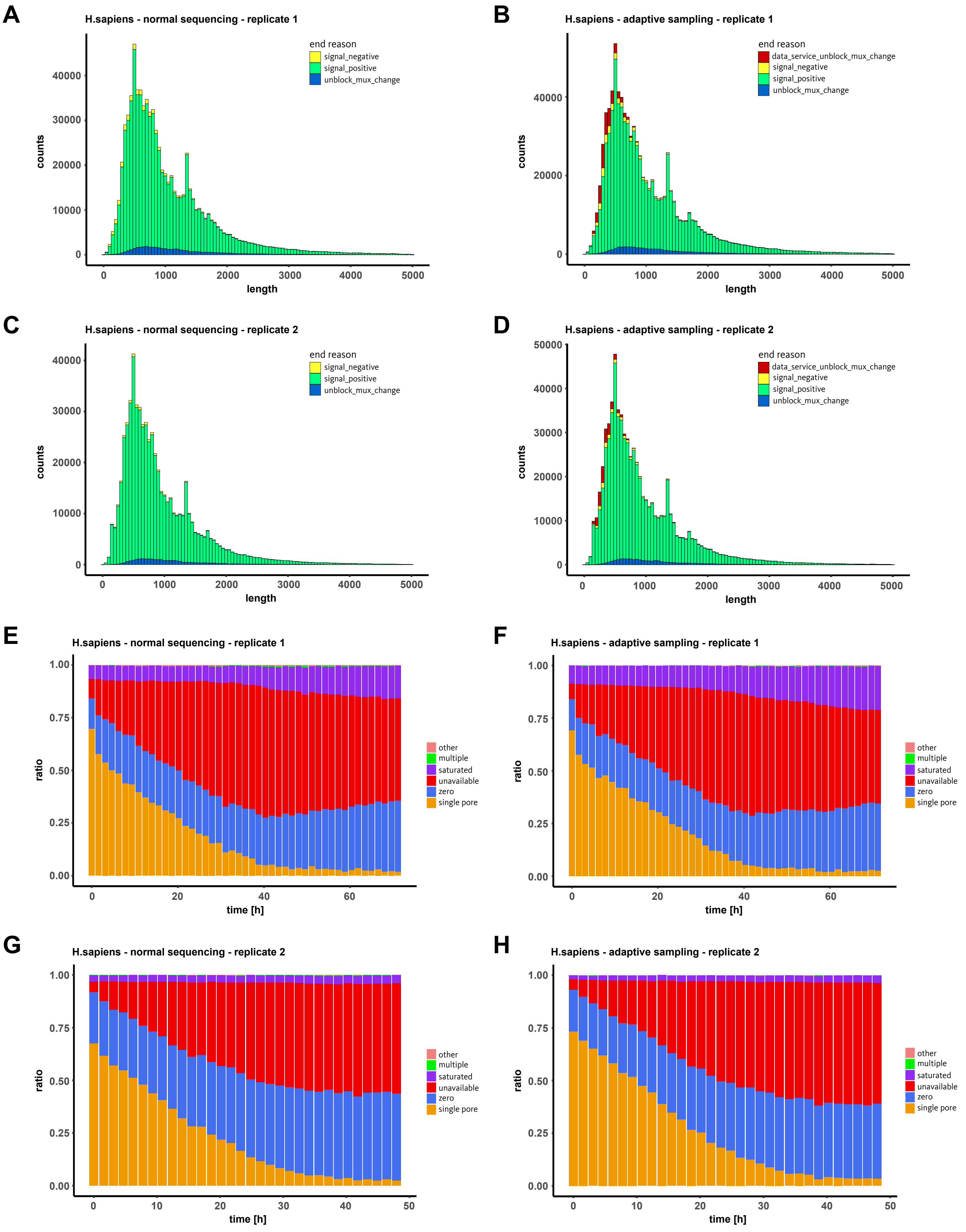

A
H.sapiens - normal sequencing - replicate 1
end reason
signal_negative
signal_positive
unblock_mux_change
counts
0
10000
20000
30000
40000
length
0
1000
2000
3000
4000
5000
B
H.sapiens - adaptive sampling - replicate 1
end reason
data_service_unblock_mux_change
signal_negative
signal_positive
unblock_mux_change
counts
0
20000
40000
length
0
1000
2000
3000
4000
5000
C
H.sapiens - normal sequencing - replicate 2
end reason
signal_negative
signal_positive
unblock_mux_change
counts
0
10000
20000
30000
40000
length
0
1000
2000
3000
4000
5000
D
H.sapiens - adaptive sampling - replicate 2
end reason
data_service_unblock_mux_change
signal_negative
signal_positive
unblock_mux_change
counts
0
10000
20000
30000
40000
50000
length
0
1000
2000
3000
4000
5000
E
H.sapiens - normal sequencing - replicate 1
ratio
0.00
0.25
0.50
0.75
1.00
time [h]
0
20
40
60
other
multiple
saturated
unavailable
zero
single pore
F
H.sapiens - adaptive sampling - replicate 1
ratio
0.00
0.25
0.50
0.75
1.00
time [h]
0
20
40
60
other
multiple
saturated
unavailable
zero
single pore
G
H.sapiens - normal sequencing - replicate 2
ratio
0.00
0.25
0.50
0.75
1.00
time [h]
0
10
20
30
40
50
other
multiple
saturated
unavailable
zero
single pore
H
H.sapiens - adaptive sampling - replicate 2
ratio
0.00
0.25
0.50
0.75
1.00
time [h]
0
10
20
30
40
50
other
multiple
saturated
unavailable
zero
single pore

### Figure S6

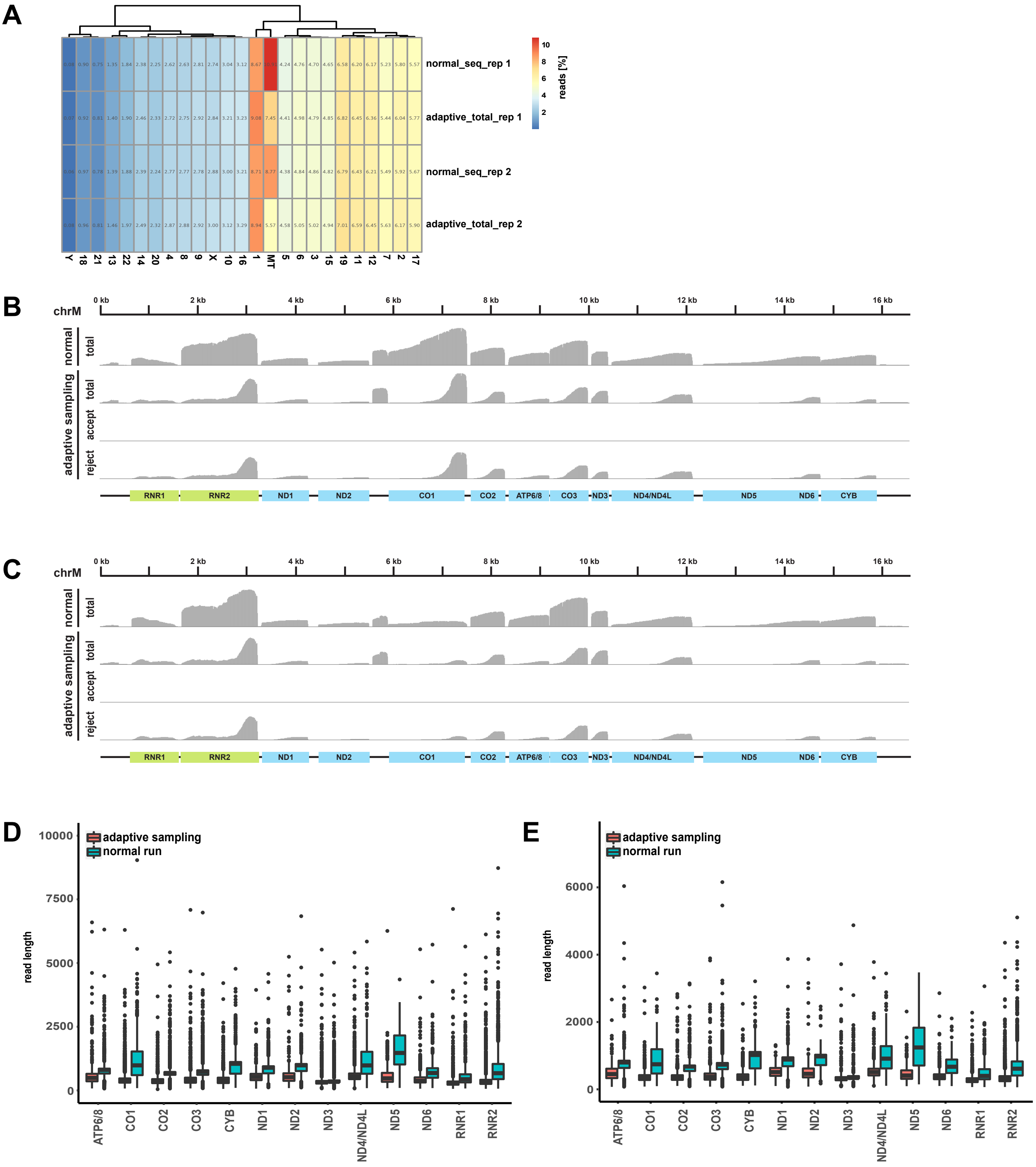
A
normal_seq_rep 1
adaptive_total_rep 1
normal_seq_rep 2
adaptive_total_rep 2
reads [%]
Y 18 21 13 22 14 20 4 8 9 X 10 16 1 MT 5 6 3 15 19 11 12 7 2 17
B
chrM
0 kb
2 kb
4 kb
6 kb
8 kb
10 kb
12 kb
14 kb
16 kb
normal
total
adaptive sampling
total
accept
reject
RNR1
RNR2
ND1
ND2
CO1
CO2
ATP6/8
CO3
ND3
ND4/ND4L
ND5
ND6
CYB
C
chrM
normal
total
adaptive sampling
total
accept
reject
D
adaptive sampling
normal run
read length
ATP6/8 CO1 CO2 CO3 CYB ND1 ND2 ND3 ND4/ND4L ND5 ND6 RNR1 RNR2
E
adaptive sampling
normal run
read length
ATP6/8 CO1 CO2 CO3 CYB ND1 ND2 ND3 ND4/ND4L ND5 ND6 RNR1 RNR2

### Figure S7

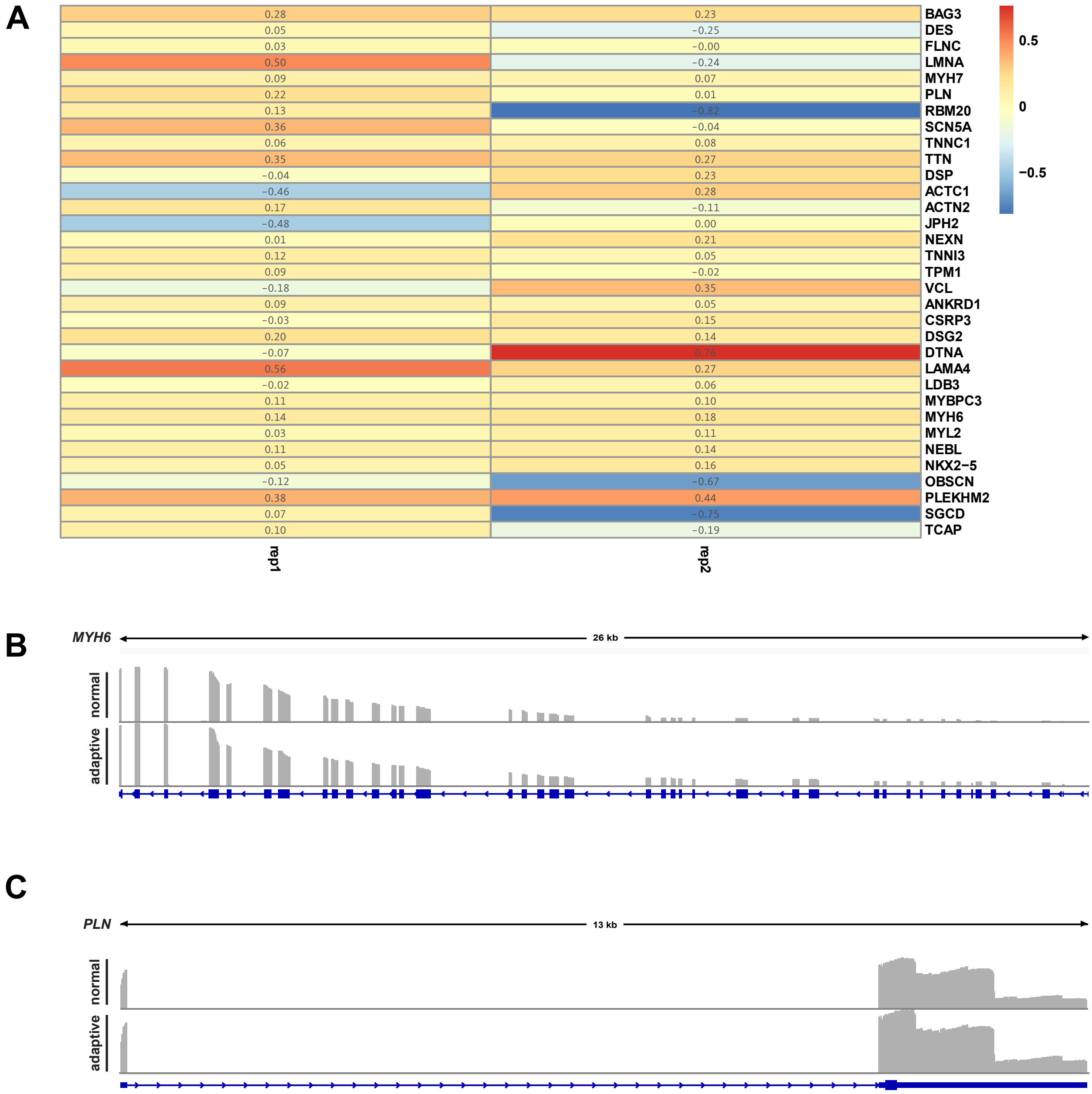
A
0.28 0.23 BAG3
0.05 −0.25 DES
0.03 −0.00 FLNC
0.50 −0.24 LMNA
0.09 0.07 MYH7
0.22 0.01 PLN
0.13 −0.82 RBM20
0.36 −0.04 SCN5A
0.06 0.08 TNNC1
0.35 0.27 TTN
−0.04 0.23 DSP
−0.46 0.28 ACTC1
0.17 −0.11 ACTN2
−0.48 0.00 JPH2
0.01 0.21 NEXN
0.12 0.05 TNNI3
0.09 −0.02 TPM1
−0.18 0.35 VCL
0.09 0.05 ANKRD1
−0.03 0.15 CSRP3
0.20 0.14 DSG2
−0.07 0.76 DTNA
0.56 0.27 LAMA4
−0.02 0.06 LDB3
0.11 0.10 MYBPC3
0.14 0.18 MYH6
0.03 0.11 MYL2
0.11 0.14 NEBL
0.05 0.16 NKX2−5
−0.12 −0.67 OBSCN
0.38 0.44 PLEKHM2
0.07 −0.75 SGCD
0.10 −0.19 TCAP
rep1
rep2
0.5
0
−0.5
B
MYH6
26 kb
normal
adaptive
C
PLN
13 kb
normal
adaptive

| chromosome | normal_shamRV | as_all_shamRV | as_unblocked_shamRV | normal_dep2 | as_all_dep2 | as_unblocked_dep2 |
|---|---|---|---|---|---|---|
| 1 | 89168 | 63948 | 43964 | 71998 | 60462 | 38437 |
| 2 | 22118 | 21888 | 150 | 19296 | 21948 | 82 |
| 3 | 11746 | 11497 | 45 | 10323 | 11697 | 28 |
| 4 | 14652 | 13933 | 42 | 13235 | 14995 | 19 |
| 5 | 22970 | 22264 | 59 | 19688 | 22338 | 32 |
| 6 | 12711 | 12524 | 37 | 10839 | 12557 | 14 |
| 7 | 27916 | 27498 | 79 | 23051 | 26049 | 49 |
| 8 | 14834 | 14486 | 35 | 13125 | 14911 | 24 |
| 9 | 20744 | 20147 | 57 | 17905 | 20728 | 31 |
| 10 | 16642 | 15804 | 49 | 14153 | 15822 | 29 |
| 11 | 22724 | 22371 | 58 | 19192 | 22296 | 39 |
| 12 | 7047 | 6890 | 21 | 5967 | 6690 | 13 |
| 13 | 9188 | 8914 | 26 | 7844 | 8844 | 18 |
| 14 | 9624 | 9261 | 22 | 8139 | 9750 | 18 |
| 15 | 16710 | 16467 | 57 | 14888 | 17240 | 32 |
| 16 | 6508 | 6507 | 19 | 5946 | 6582 | 10 |
| 17 | 10753 | 10547 | 34 | 9186 | 10798 | 17 |
| 18 | 5489 | 5434 | 14 | 4757 | 5473 | 8 |
| 19 | 8571 | 8087 | 25 | 7471 | 8248 | 13 |
| X | 7248 | 7131 | 28 | 6212 | 7195 | 10 |
| Y | 79 | 66 | 0 | 58 | 81 | 0 |
| MT | 183797 | 97803 | 84727 | 132385 | 90615 | 74960 |

| Chromosome | normal_rep1 | as_all_rep1 | as_unblocked_rep1 | normal_rep2 | as_all_rep2 | as_unblocked_rep2 |
|---|---|---|---|---|---|---|
| 1 | 65032 | 74754 | 390 | 49029 | 56737 | 99 |
| 2 | 43499 | 49721 | 51 | 33321 | 39148 | 20 |
| 3 | 35251 | 39466 | 54 | 27350 | 31872 | 12 |
| 4 | 19633 | 22439 | 23 | 15615 | 18219 | 8 |
| 5 | 31797 | 36314 | 49 | 24673 | 29104 | 7 |
| 6 | 35693 | 40987 | 33 | 27250 | 32038 | 14 |
| 7 | 39220 | 44829 | 42 | 30898 | 35733 | 9 |
| 8 | 19745 | 22625 | 20 | 15606 | 18306 | 7 |
| 9 | 21060 | 24080 | 26 | 15626 | 18571 | 8 |
| 10 | 22815 | 26415 | 25 | 16898 | 19813 | 9 |
| 11 | 46494 | 53126 | 54 | 36181 | 41826 | 21 |
| 12 | 46273 | 52386 | 65 | 34953 | 40947 | 17 |
| 13 | 10096 | 11534 | 35 | 7801 | 9271 | 18 |
| 14 | 17863 | 20292 | 23 | 13480 | 15838 | 8 |
| 15 | 34870 | 39985 | 48 | 27121 | 31378 | 16 |
| 16 | 23384 | 26622 | 29 | 18059 | 20894 | 6 |
| 17 | 41750 | 47533 | 46 | 31935 | 37464 | 11 |
| 18 | 6720 | 7589 | 5 | 5456 | 6090 | 6 |
| 19 | 49363 | 56154 | 59 | 38224 | 44512 | 18 |
| 20 | 16867 | 19193 | 22 | 12584 | 14700 | 3 |
| 21 | 5619 | 6664 | 7 | 4419 | 5122 | 0 |
| 22 | 13782 | 15624 | 15 | 10584 | 12477 | 7 |
| X | 20534 | 23374 | 29 | 16234 | 19037 | 7 |
| Y | 570 | 547 | 1 | 359 | 479 | 0 |
| MT | 81783 | 61355 | 44396 | 49382 | 35354 | 25332 |
